# Partitioning the roots of interactions between microbes (PRISM): environment-supplied resources versus species-produced mediators

**DOI:** 10.64898/2026.05.01.721678

**Authors:** Vaishnavi Warrier, Babak Momeni

## Abstract

Microbial species in a community can interact through competition for resources present in the environment, as well as inhibition or facilitation by metabolites produced by other species. Understanding the relative contribution of these two types of interaction will help us modulate bacterial interactions more effectively. Our work focuses on partitioning the impact of metabolites that mediate the interactions between bacteria into the contribution of resources present in the environment versus that of mediators released as by-products of cellular activities. For this, we create a range of conditions in which the ratio of environmentally supplied resources (R) and the species-produced metabolites (M) is modulated to infer the contribution of each to the overall interaction. We performed this assay with six different nasal bacterial strains and saw an array of outcomes in terms of how secreted metabolites by one strain affected other nasal strains. Metabolites produced by *Staphylococcus* suppressed the growth of most strains tested. In contrast, metabolites produced by commensal strains such as *Corynebacterium accolens* and *Corynebacterium tuberculostearicum* could aid the growth of *Staphylococcus aureus* and *Staphylococcus epidermidis* strains. Interestingly, the growth of *Staphylococcus epidermidis* benefited from metabolites produced by four out of the five strains tested. Our proposed assay offers additional insights into the roots of bacterial interactions, enabling applications such as engrafting helpful probiotics and knocking out harmful bacteria.

## Introduction

Microbe-microbe interactions play an important role in determining the bacterial communities’ structure and function. These interactions include the exchange of metabolites that support species’ growth and establishment. These exchanges, also referred to as cross-feeding, allow microbes to exploit resources more efficiently and can be critical for the formation and maintenance of microbial communities [1]. Interactions between microbes can be tightly linked to community dynamics [2]. By better understanding interactions, we can manipulate communities, influencing outcomes in areas like disease treatment and microbiome engineering [3][4][5].

Microbial interactions have diverse mechanisms. Microbes may influence each other, for example, by producing toxins or by metabolic cross-feeding. Microbes can also directly or indirectly compete with each other for limited resources like space and nutrients. A combination of these interactions makes determining how species interact with one another a challenge in diverse ecosystems. Further complicating the situation, microbial interactions can depend on the abiotic environment it is present in as well as its biotic context [6].

Past efforts on characterizing microbial interactions offer us a myriad of culture-dependent methods [7]. These methods, combined with sequencing and metabolomics, can provide a detailed understanding of the mechanisms behind these interactions [8]. In addition to these extensive culture-based approaches of identifying bacterial interactions, several mathematical modeling frameworks exist for predicting the dynamics of ecological communities. One of the most common modeling techniques used to study this is the generalized Lotka-Volterra model [9]. The LV model captures how one microbe stimulates/inhibits the growth of another microbe based on intrinsic growth parameters and pairwise interactions. However, it does not take into consideration the context or the mechanism of interactions [10][11]. For example, a negative interaction coefficient in gLV might represent two species interacting neutrally while competing for a nutrient or it might represent an inhibitory toxin with little or no competition for shared resources. Apart from gLV, other models have also been developed with focus on certain types of interactions. The Consumer-Resource model or a full metabolic model, for example, emphasize competition for shared resources and cross-feeding [12], still only partially capturing the complexity of interactions among microbes at the expense of more model complexity.

Past reports have used empirical and modeling approaches to examine the contribution of competition to microbial interactions. A study by Huang et al., has developed a framework to estimate multispecies niche overlap of 15 human gut commensals combining their individual metabolomics data, growth measurements in spent media, and mathematical modeling to study the effect of resource competition in pairwise interactions [13]. Another such study performed by Wagner and colleagues used genome-scale metabolic reconstructions of 250 community assembly sequences. They showed that nutrient competition among residents in these communities strengthens community resistance to invaders while cross-feeding interactions can weaken that resistance [14]. Although these studies have simultaneously studied the effect of resources and metabolic mediators, they have not teased apart these effects in each interaction.

To get an in-depth understanding of the nature of microbial interactions, it would be helpful to tease apart the effect of environment-supplied resources (R) from that of species-produced mediators (M). Some of these mediators produced from species, such as short chain fatty acids, lipopolysaccharides, extracellular vesicles, enzymes, and vitamin by-products, can have potential health benefits in modulating the microbiome and supporting immune functions. These molecules, often referred to as postbiotics, are more stable and effective than live probiotics [15]. Discovery of postbiotics using methods like chromatography and mass-spectrometry can often be expensive and time consuming due to a lack of standardized screening protocol in place [16]. Here we propose a simple assay that provides insights into whether species-produced compounds are mediating the interactions between microbes. In addition to offering clues about interaction mechanisms, these insights help decide if a certain interaction is worthy of being subjected to more elaborate chromatography and mass-spectrometry methods.

## Results

### A linear model can combine the contribution of resources and mediators

In the simplest form, the contribution of environment-supplied resources versus species-produced mediators can be considered to be linearly additive. We use a linear model to represent the effects of resources and mediators on microbial growth, by assigning coefficients of dependencies to each variable.

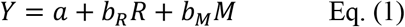

Where, *Y* is the carrying capacity of a species exposed to a partner, a is the intercept (constant term), *b*_*R*_ represents the dependency on *R* (i.e. resource dependence), *b*_*M*_ represents the dependency on *M* (i.e. the effect of partner-produced mediators) and *R* and *M* are the independent variables representing the resources and mediators, respectively.

When examining a microbe (species 1) growing in the supernatant of a partner (species 2), the above linear model, is influenced by two variables. The value *R* represents the resource dependence of species 1 on the nutrients or compounds present in the supernatant of species 2 whereas the value *M* represents the effect of mediators present in the supernatant of species 2 on growth of species 1. The effect of *R* on the growth of species 1 is captured by the coefficient *b*_*R*_. If *b*_*R*_ is positive, providing more resources leads to more growth of species 1, whereas if *b*_*R*_ is negative, more supply of shared environmental resources causes a stagnancy in the growth of that species. Similarly, the effect of *M* (produced by species 2) on the growth of species 1 is captured by *b*_*M*_. A positive *b*_*M*_ indicates that compounds produced by the partner favor more growth of species 1, with mechanisms that may include favorable signaling molecules or beneficial metabolic by-products. A negative *b*_*M*_, on the other hand, indicate the presence of growth inhibiting interaction mechanisms that suppress the growth of species 1, for example by releasing inhibitors or other anti-microbials. Overall, interactions are determined by both the availability of resources and the additional effects mediated by components of the supernatant, with coefficients *b*_*R*_ and *b*_*M*_ representing the strength and direction of these influences. The intercept, a, represents the baseline bacterial growth when both *R* and *M* are zero. In most cases, we see that the value of a is very close to 0, suggesting that the species being examined shows no growth in the absence of *R* and *M*.

### Correction terms can account for the residual resources in conditioned media

To predict the effect of supernatant and its partitioning, in Eq. (1) we assume that the supernatant obtained from a species is completely spent, such that the same bacteria can no longer grow in that supernatant. However, this assumption does not always hold. Often the filtrate obtained after growth of a bacterium is conditioned medium, rather than a fully spent medium, and can still support the growth of the same bacteria. This leads to error in estimated interactions since the values of *R* and *M* will not be a true reflection of resources/mediators in the spent media. To circumvent this error, we correct for the residual resources in the conditioned medium by introducing a variable *f* which is the ratio of carrying capacity of conditioned medium to that of the fresh medium (i.e. *K*_*C*_ */K*_*R*_). We use a simple set of equations to derive these values.

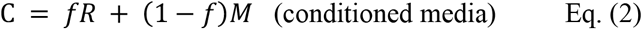

*f* is the fraction of *R* in the mixture, while (1-*f*) is the fraction of *M*. This equation models how the conditioned media (C) is a blend of the spent and non-spent components in the media controlling the relative amounts of each.

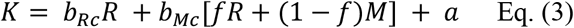

Combining the two equations above, we get a new equation for carrying capacity where K is a combination of the effects of F (directly) and the mixture of F and S (through the fraction *f*).

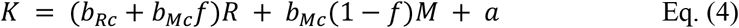

Above equation is obtained after further expanding K from equation 3.

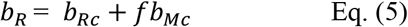

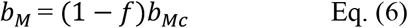

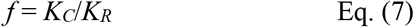

Thus, using this *f* value in Eqs. (5)-(7), we can account for the residual resources in the conditioned medium and obtain a more accurate representation of the contributions of *M* (i.e. spent medium) and *R* (i.e. fresh medium) to the growth of the focal strain.

### Assessing the growth of species in the supernatant of other species gives us an overall interaction coefficient but not the mechanism behind it

Before we look at the partitioned interactions between spent medium of different strains, we first examined the overall effects of the spent medium. For this, we examined the growth of species in the supernatants obtained from five different nasal bacteria [10]. We compared a diluted rich culture medium, MOPS-buffered 10% Todd-Hewitt broth with yeast extract supplemented with Tween80 (10%THY+T, Figure 1B) with a defined medium, Baseline Amino Acid Defined supplemented with Tween80 (BAAD+T, Figure 1A) [10]. The bacterial growth in fresh and spent media was probed by measuring the carrying capacity in these different conditions after 24 hours.

**Figure 1.**
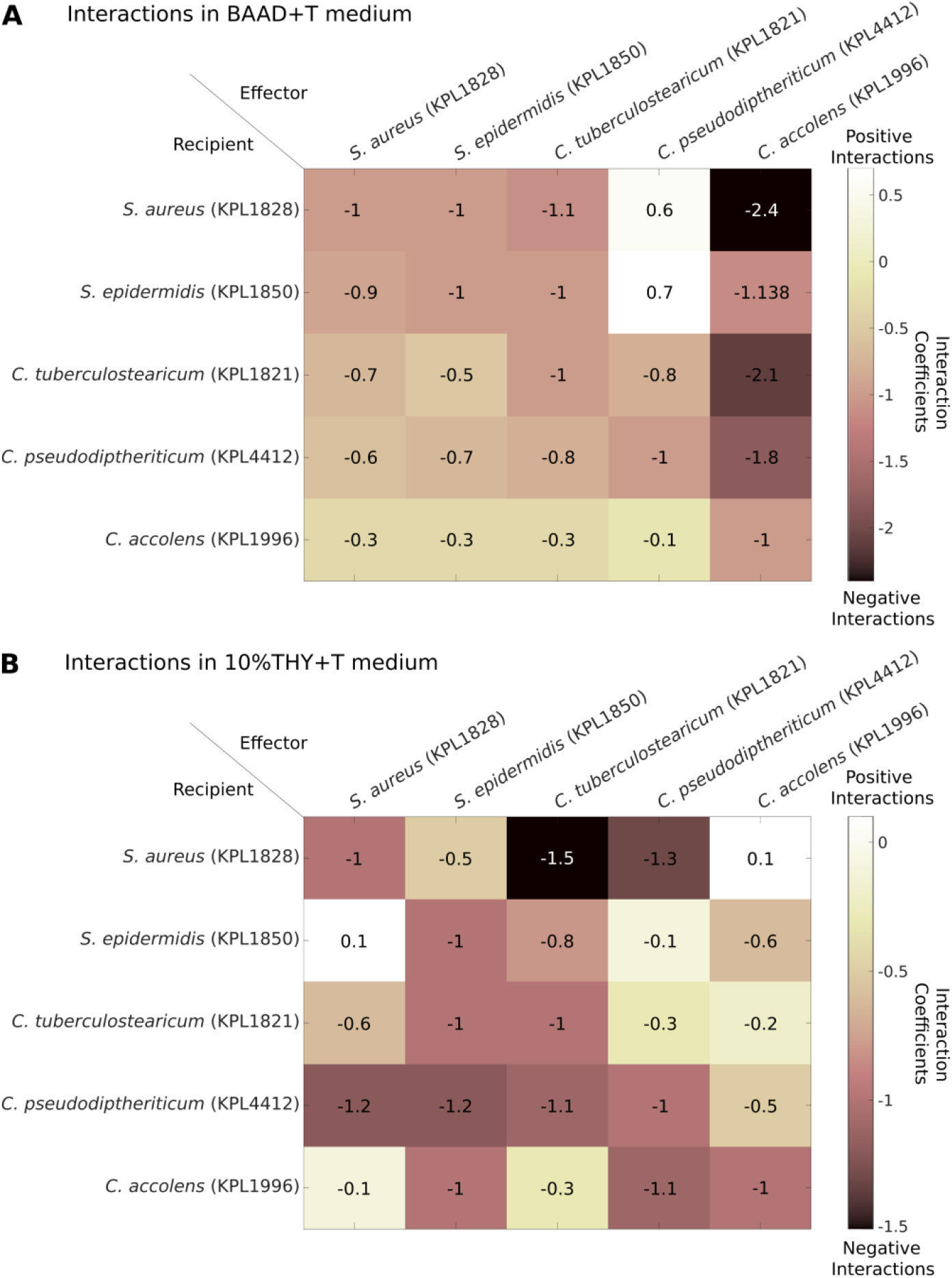
Interactions among nasal bacteria inferred from corrected supernatant assay depend on the growth medium. The interaction coefficients are calculated by measuring the carrying capacity of the recipient species in the supernatant of the effector species. We used the correction described in Supplemental Material S.1 and Eq. (8) to correct the effect that the supernatant may not be fully spent. (A) The interaction coefficients for five nasal strains in the supernatant of all the other strains grown in BAAD+T medium are shown. These interactions were mostly negative, with the exception of *Staphylococcus* strains in *C. pseudodiptheriticum* supernatant. (B) The interaction coefficients for five nasal strains in the supernatant of all the other strains grown in 10%THY+T medium are shown. These interactions were more diverse in nature with some highly negative interactions (*S. aureus* in *C. tuberculostearicum*) as well as positive interactions (*S. aureus* in *C. accolens* and *S. epidermidis* in *S. aureus*).

When calculating the interactions coefficients (*c*_*ij*_; see Supplemental Material S.1), we noted that the assumption that supernatants are cell-free spent media (CFSM) led to inconsistent results across different trials and self-interaction terms (*c*_*ii*_) were different from the predicted value of −1 (expected from logistic growth). We hypothesized that this difference is because the supernatant or cell-free conditioned media (CFCM) were not fully depleted by the effector strains. We used the growth of each species in its own supernatant to correct for this effect (see the derivation in Supplemental Material S.1). With this correction, we find that the interaction coefficients can be more accurately calculated as

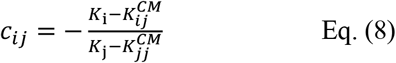

where *K*_i_ and *K*_j_ are the carrying capacity of species *i* and *j* in a monoculture, respectively, and 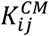 and 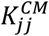 are the carrying capacity of species *i* and *j* in the CFCM of species *j*, respectively.

Based on the corrected values of the interaction coefficients and comparing the two different media, we see an overall negative, but not large, interaction coefficients between nasal isolates (Figure 1). Interactions are more prominent in BAAD+T, with larger negative interaction coefficients. In 10%THY+T medium, interaction coefficients have a wider distribution between the range of positive versus negative interactions. These results give us a basic picture of how different nasal bacteria interact, but they do not tell us if interactions are due to competition for shared resources or if species have produced byproducts that influence other species. For example, an interaction coefficient of −0.8 could mean that there is 80% niche overlap and thus resources that support the recipient species are remained (at 20% of what the fresh medium provides) in the supernatant of the effector strain. However, the same interaction coefficient could also mean a 60% niche overlap (40% leftover nutrients) along with a species-produced byproduct in the supernatant that support an additional 20% growth, or a 100% niche overlap (0% leftover nutrients) along with a species-produced toxin in the supernatant that reduces growth by 20%. The supernatant assay is not capable of distinguishing these cases from each other.

### Identify the contribution of resources versus mediators by examining the growth of a focal strain exposed to different doses of another strain’s cell-free conditioned medium

To identify the contribution of resources versus mediators, instead of examining only the effect of conditioned media, we mixed the conditioned media with fresh media (or a concentrated or diluted version of it) at different fractions (Figure 2). For each strain-CFCM pair, we mixed seven different concentrations of CFCM and 3 different concentrations of the baseline medium. The concentrations of the fresh medium were chosen within the range that the growth rate and carrying capacity had a linear relation [10] to make it easier to linearly decompose their effects (Supplemental Material S.2). We then measured the growth dynamics and calculate the carrying capacity in all the conditions over 24 hours (OD_600_ measured every 10 minutes).

**Figure 2.**
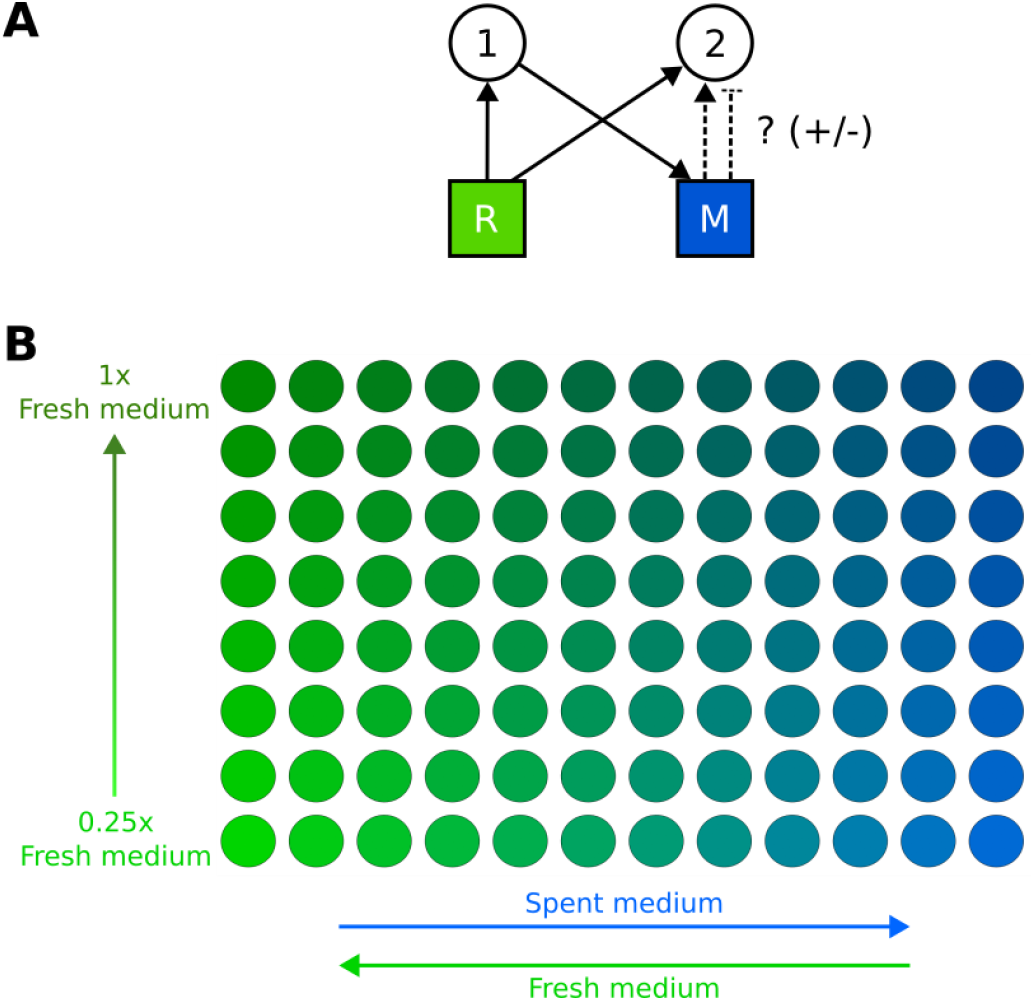
The outline of PRISM assay for inferring the contribution of resources versus mediators is shown. (A) This conceptual figure shows the interaction between two species in the presence of a shared resource (R) and a mediator (M) released into the environment by species 1. While species 1 consumes the available resources for its own growth, it can also release mediators that have a positive or negative effect on species 2. Thus, the growth of species 2 not only depends upon the resource competition for the leftover nutrients, but also the chemical environment modified by species 1. (B) The assay to monitor the growth of species 2 in the conditioned medium of species 1 is shown. This assay includes a range of conditions in which the ratio of environmentally supplied resources (R) (the fresh medium) and the species-produced metabolites (M) (the conditioned medium produced by species 1) is modulated to infer the contribution of each to the overall interaction towards species 2.

### Staphylococcus aureus inhibits the growth of other nasal commensal bacteria

We conducted our PRISM assay on five different representative nasal strains to better understand the nature of their interactions. All the five strains showed a strongly positive growth on fresh media, which means that these strains grow well when there is adequate supply of nutrients in this growth culture. The results showed that for most strains an inhibitory mediator against them was present in the conditioned media of *S. aureus*. Although this trend was consistent among all the strains, the negative effects of *S. aureus* seemed more prominent in the chemically defined BAAD+T medium. The reason could be that *S. aureus* produces more inhibitory compounds in a defined media or the recipient strains may be more susceptible to *S. aureus*-produced inhibitory compounds in such an environment. On the other hand, in a more complex medium such as 10%THY+T, we see less inhibitory effect of the supernatant. Again, this could be a result of lowered toxin expression in an environment with less metabolic stress or the recipient strains might be more protected by some of the nutrients present in that environment.

The graphs shown in Figure 3 characterizes the linear model equation (*Y* = *a* +*b*_*R*_ *R* +*b*_*M*_ *M*) for dependencies of the growth of different strains in the fresh medium (*b*_*R*_), or the dependency on supernatant of *S. aureus* (*b*_*M*_). Total growth (here, based on maximum measured OD_600_) is modeled as a combination of the contribution by R (represented as *b*_*R*_*R*) and the contribution by M (represented as *b*_*M*_ *M*).

**Figure 3.**
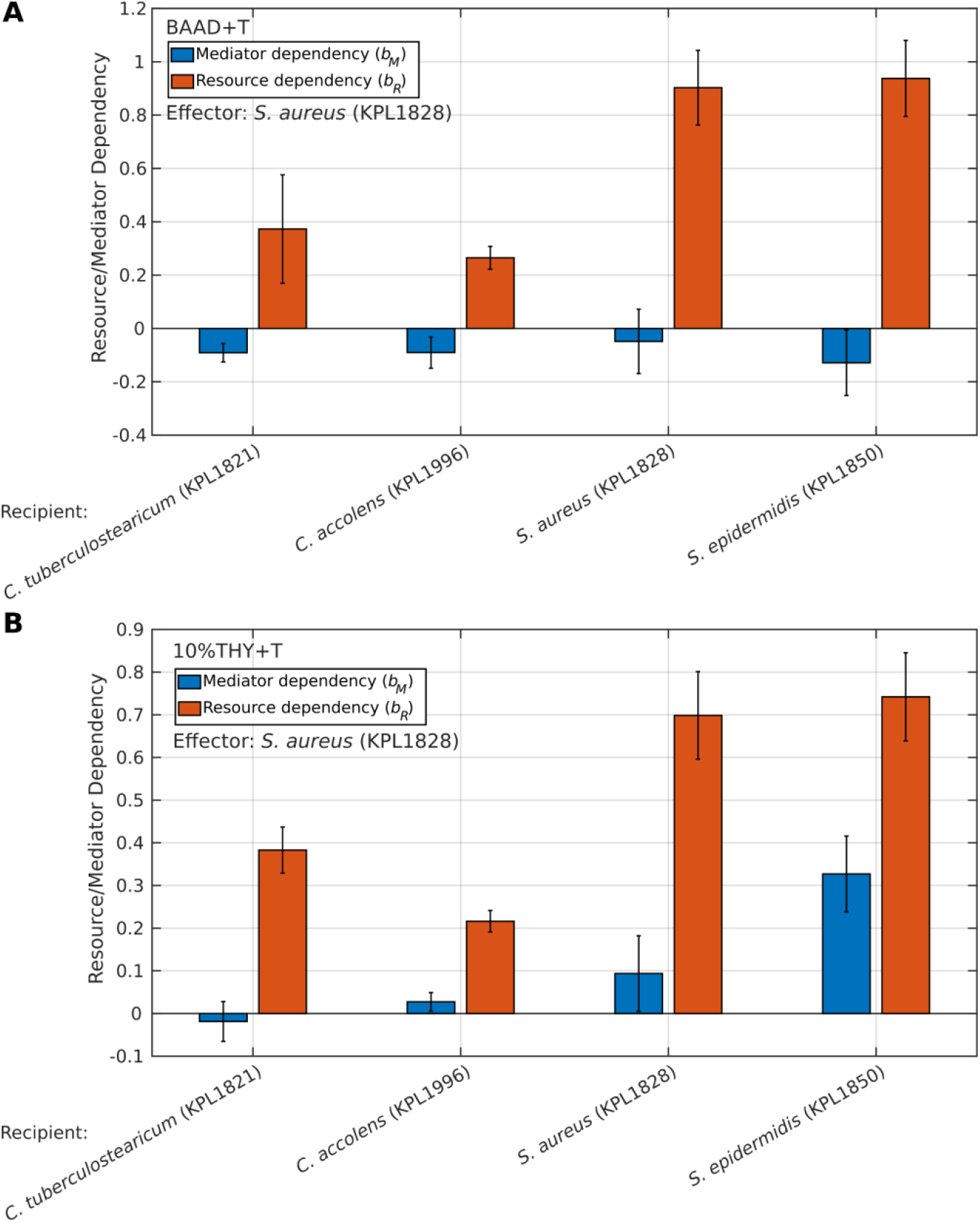
Estimating the contribution of environment-supplied resources versus species-produced mediators for five different nasal strains. This bar graph figure compares the growth changes of four different nasal bacteria strains under two different conditions. (A) The effect of BAAD+T cultured *S. aureus* supernatant versus fresh medium on four nasal strains. (B) The effect of 10%THY+T cultured *S. aureus* supernatant versus fresh medium on four nasal strains. The orange bars represent the dependency of growth on fresh media or supplied resources and the blue bars represent the dependency of the growth of strains on the supernatant of *S. aureus*. The dependency of the species to grow either in the supernatant or fresh medium is calculated by measuring the carrying capacity per unit medium concentration. Error bars show the 95% confidence interval obtained from the linear regression of slopes.

### Corynebacterium-produced mediators enhance the growth of Staphylococcus strains

To assess the impact of *Corynebacterium* effect on other nasal strains, we looked at the growth of multiple strains in fresh media and *Corynebacterium* conditioned media (Figure 4).

**Figure 4.**
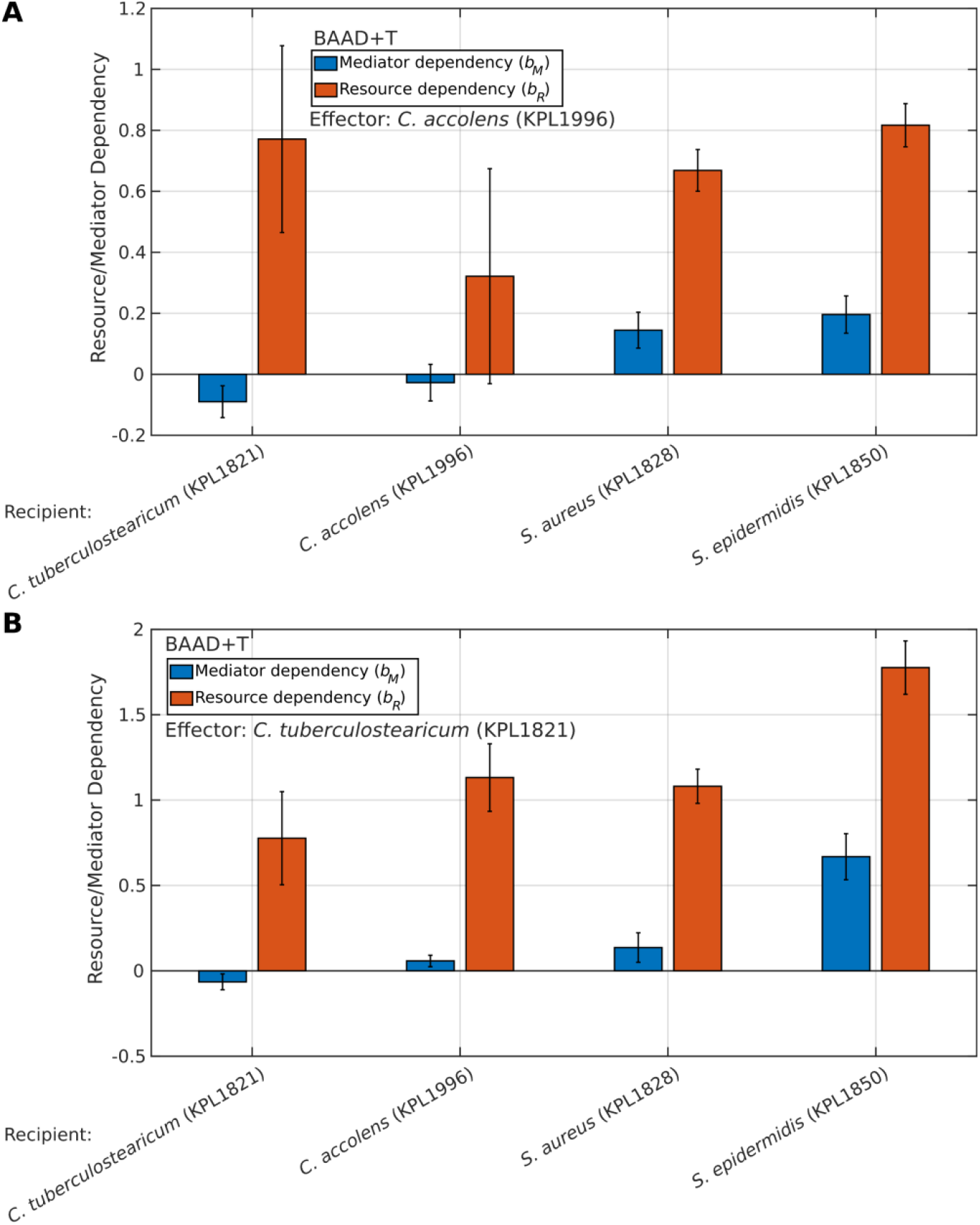
*Corynebacterium* species appear to produce mediators that support the growth of Staphylococcus species. These bar graph figures show the growth changes of four different nasal bacteria strains under the effect of two different *Corynebacterium* strains. (A) The effect of BAAD+T cultured *C. accolens* supernatant versus fresh medium on four nasal strains. (B) The effect of BAAD+T cultured *C. tuberculostearicum* supernatant versus fresh medium dependency of the five strains. The orange bars represent the dependency of growth on fresh media or supplied resources and the blue bars represent the dependency of the growth of strains on the supernatant of *C. accolens* and *C. tuberculostearicum* respectively. The dependency of the species when growing either in the supernatant or fresh medium is calculated by measuring the carrying capacity per unit medium concentration. Error bars show the 95% confidence interval obtained from the linear regression of slopes.

**Figure 5.**
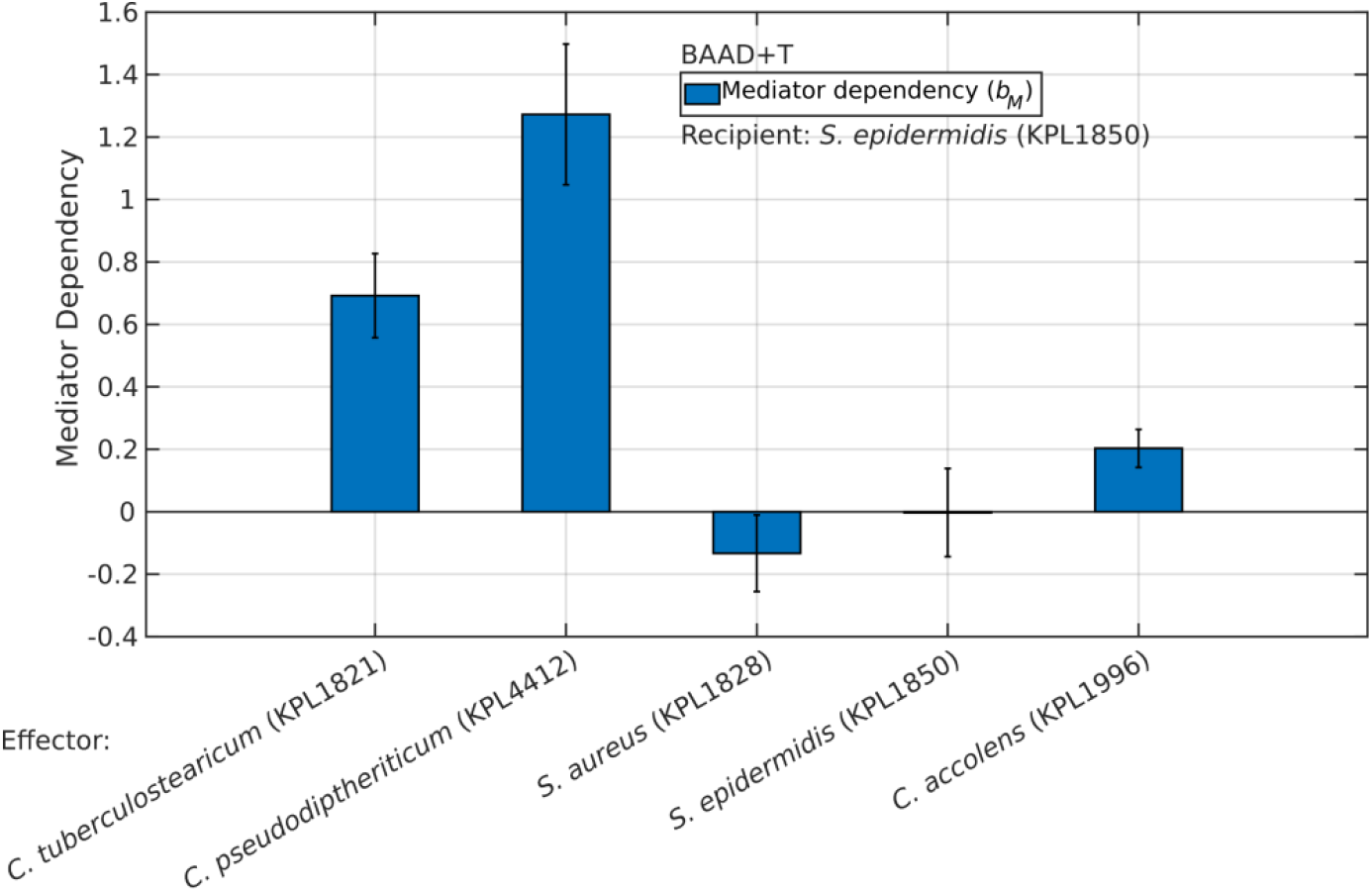
Growth of *S. epidermidis* is improved by the species-produced mediators of most other nasal strains. This bar graph shows the growth changes of *S. epidermidis* in different nasal bacteria supernatants. The blue bars represent the dependency of the growth of *S. epidermidis* in the supernatant of the respective strains. The dependency of the species to grow in the supernatant is calculated by measuring the carrying capacity per unit concentration. Error bars show the 95% confidence interval obtained from the linear regression of slopes.

Our results suggest that the effect of the supernatants of *Corynebacterium* species depends on the recipient strain, promoting the growth of some strains and inhibiting to the growth of other strains. These interactions reveal a highly species-specific effect of *Corynebacterium* strains on the other nasal strains in the environment. Of note, the growth of *S. epidermidis* enhanced by the species-produced mediators of both *C. accolens* and *C. tuberculostearicum*. These effects could be a result of *Corynebacterium* releasing useful metabolites that directly enhance *S. epidermidis* growth or by eliminating inhibitory compounds indirectly aiding the growth of *S. epidermidis*. It is interesting to see such a preferential aid in growth of *S. epidermidis* considering it is a beneficial commensal in the nasal microbiome.

We also saw a positive effect of *Corynebacterium-* produced mediators on *S. aureus*, but this effect was not as prominent as the effect on *S. epidermidis*. Taken together, we see that *Corynebacterium*-produced mediators promoted the growth of *Staphylococcus* strains among the nasal bacteria that we studied.

### S. epidermidis benefits from mediators produced by most other nasal bacteria

We observed that *S. epidermidis* experienced enhanced growth when exposed to mediators produced by other nasal commensal strains. This positive effect was the highest from *C. pseudodiptheriticum, C. tuberculostearicum* and *C. accolens*. While a variety of factors could be responsible for such a trend, a possible scenario is that *S. epidermidis* could readily use a common byproduct of bacterial metabolism that gets released by the other commensals.

## Discussion

We propose a simple linear model and a corresponding PRISM assay to distinguish between partner produced mediators and resource competition in the interaction between microbes. We demonstrate that interactions between different bacteria are not merely governed by competition for resources but also by byproducts released by microbes during their growth. Using a group of nasal bacterial strains, we examine how their interactions depended on competition versus other species-produced mediators. We find that *S. aureus* produces mediators that are inhibitory to many other nasal bacteria. In contrast, mediators produced by *Corynebacterium* strains appear to be beneficial to *Staphylococcus* strains. Notably, *S. epidermidis* appears to benefit from mediators produced by most other nasal bacterial strains, pointing to its possible role as a generalist consumer of byproducts in the nasal microbiota.

The linear model of resources and mediators (Eq (1) and Fig. 2) provides a simplified yet powerful method for compartmentalizing the contribution of resources and mediators in microbial interactions with possible practical implications [17], [18], [19]. This model lets us quantitatively partition between resources and mediators with an easy fitting mechanism based on experimentally measured growth properties such as carrying capacities. The model can be used validly under conditions tested within the linear ranges of growth responses. Even though in principle three independent data points are adequate to parametrize the model, we typically use more conditions (21 in our standard implementation of the PRISM assay) to test for linearity and gain more confidence in estimated slopes of the linear model.

The distinction between competition for resources versus species-produced mediators has profound implications in our understanding of bacterial interactions from an ecological perspective and for devising intervention strategies to manipulate bacterial communities. Growth-promoting mediators can be identified and utilized for favoring beneficial commensals as a prebiotic strategy to restructure microbiota. The mediators promoting growth factors may represent several cross-feeding pathways stabilizing co-existence and community resilience.

These insights provide a framework for predicting community dynamics. Similarly, inhibitory mediators (or their analogs) can be used to specifically diminish the growth of harmful pathogens as narrow-spectrum antimicrobials. The PRISM assay helps target potential candidates for interaction mediators to be further investigated to identify molecular mechanisms of interactions. Overall, our work helps to better understand the innerworkings of communities as a combination of competition for shared resources along with interactions via species-produced effects.

By demonstrating the assay using a known antibiotic (Supplemental Material S.3 and Figure S1), we also open doors to another approach for the discovery of antimicrobial compounds from supernatant. Standard antibiotic testing assays test purified compounds against a strain of interest providing limited information on the ecological bearing on the other strains in that environment. The PRISM assay removes the requirement for compound isolation by directly inferring the effect of compounds produced by one species on the growth of strains of interest.

The PRISM assay offers insights about the magnitude and nature of interactions within a community as highlighted by our investigation of nasal bacteria. We see that *S. aureus* may be behaving as a dominant competitor exerting negative effects on the growth of other strains. This inhibitory effect is stronger in a chemically defined BAAD+T medium, which suggests an important environment-dependent factor in determining interbacterial interactions. In contrast, in 10%THY+T there is a possibility of a reduced toxin expression and/or protective effects of surplus nutrients. These findings reinforce some of the previous findings; for example, a study by Peschel *et al*., has shown that limiting nutrients in the nasal environment shifts the regulation of the methionine biosynthesis pathway, offering an antimicrobial target [20]. Additionally, such conditions may lead to the activation of the quorum sensing systems in *S. aureus*, releasing phenol soluble modulins (PSMs) that have antimicrobial activities against other nasal commensals [21]. To explore such possibilities, we performed anti-SMASH analysis [22] on the genome of *S. aureus* KPL1828 to identify any biosynthetic gene clusters encoding antimicrobial peptides. However, this analysis did not find any obvious hits, possibly because many previously explored inhibitory factors (such as PSMs and δ-toxin) are small peptides that may get well represented as biosynthetic gene clusters.

Aside from having the potency to identify antimicrobials using our assay, we also have looked into the interactions amongst the other nasal commensals that can thrive as a community. We specifically focused on testing the effect of nasal *Corynebacterium* strains on other bacterial nasal strains. While we see that these strains have a mixed array of positive and negative effects on the other strains, they have a notably high positive effect on growth of the *S. epidermidis* strain. This co-habitation of *S. epidermidis* and *Corynebacterium* species has been established before in the context of human skin [23]. Previous work has also shown that skin-associated *Corynebacterium amycolatum* releases cobamides (Vitamin B12 cofactors) important for maintaining community diversity [24]; However, these trends have not been observed in nasal *Corynebacterium* species yet.

There are some intrinsic assumptions and corresponding limitations in the use of our proposed PRISM assay. First, the contributions of resources and mediators are assumed to act independently such that the model can partition their relative contributions. This assumption may not necessarily hold; however, if that is the case, the higher-order interaction term in the linear model (*b*_*RM*_*RM*) will be informative for exploring the possible nature of this mechanism. Similar to other supernatant-based assays, interactions based on direct physical contact or changes in cell properties or physiology in monocultures versus in the presence of the partner are not captured in the PRISM assay [10]. Another assumption is that the mediators present in supernatant will stay biologically active and structurally stable throughout the length of the assay. The assay also presumes that external factors such as oxygen depletion do not attribute to the effects of resources or mediators. By design, the PRISM assay does not identify the specific metabolites or pathways responsible for these effects. Additionally, since the assay is conducted under linear growth conditions, we may run the risk of oversimplifying complex nutrient environments or higher-order interactions. To simplify the interpretations, we may dilute the effects of mediators and resources to bring them to a linear response range. The scalability of interactions under such conditions requires further confirmation.

The PRISM assay developed in this paper provides a framework for disentangling the contribution of resources and metabolite effect in the pairwise interactions of bacteria. Several promising extensions could further help a wise usage of this model. One of the most apparent next steps would be to chemically characterize the role of mediator or inhibitors by applying metabolomics-based research. Further work can also be focused on accounting for higher-order interactions and non-linear responses to extend this concept to more complex multi-species communities.

## Materials and Methods

### Bacterial strains

The bacterial strains were received from the Katherine P. Lemon lab where the primary nostril isolates were isolated from the nasal vestibule of two adult volunteers under an Institutional Review Board-approved protocol. One volunteer was a *Staphylococcus aureus* carrier and one a non-carrier.

**Table 1.** Bacterial strains used in this study and their alias. The alias in this manuscript is chosen for simplicity, based on the closest phylogenetic match for each strain.

| Strain name | Alias in this manuscript |
| --- | --- |
| *Staphylococcus aureus* KPL1828 | *S. aureus* |
| *Staphylococcus* sp. KPL1850 | *S. epidermidis* |
| *Corynebacterium accolens* KPL1996 | *C. accolens* |
| *Corynebacterium tuberculostearicum* KPL1821 | *C. tuberculostearicum* |
| *Corynebacterium pseudodiphtheriticum* KPL 1989 | *C. pseudodiphtheriticum* |

### Growth conditions

To revive cells from frozen stocks, bacterial isolates were grown in 3 mL of 100% Todd Hewitt Broth + 0.5% Yeast Extract + 1% Tween80 (THY+T) until mid-exponential growth phase (typically to an OD_600_ between 0.15 and 0.3). Cells from each monoculture were then centrifuged down and re-suspended in 1× phosphate-buffered saline (PBS) before transferring a small fraction into the specific testing condition.

THY medium was made using the premixed powders of Todd Hewitt broth (BD Bacto Dehydrated Culture Media: Todd Hewitt broth, Thermo Fisher Scientific, DF0492-17-6) and yeast extract (Thermo Scientific Yeast Extract, Ultrapure, AAJ23547A1). 1% Tween-80 (Fisher BioReagents, BP338500) was added to this prepared media.

Baseline Defined Media with amino acids (BAAD+T) medium consists of 1.5 g/L of KH_2_PO_4_, 3.8 g/L of K_2_HPO_4_ (×3H_2_0), 1.3 g/L of (NH_4_)_2_SO_4_, 10 g/L of MOPS, 3 g/L of sodium citrate (×2H_2_O), 10 mL/L of the mixed vitamin stock (2 mg/L of biotin, 2 mg/L of folic acid, 10 mg/L of pyridoxine-HCl, 5 mg/L of thiamine-HCl ×2H_2_O, 5 mg/L of riboflavin, 5 mg/L of nicotinic acid, 5 mg/L of D-Ca-pantothenate, 0.1 mg/L of vitamin B12, 5 mg/L of p-aminobenzoic acid, and 5 mg/L of lipoic acid), 1 mL of SL-10 mixed trace elements stock (10 mL/L of HCl (25%; 7.7 M), 1.5 g/L of FeCl_2_ ×4H_2_O, 70 mg/L of ZnCl_2_, 0.1 g/L of MnCl_2_ ×4H_2_O, 6 mg/L of H_3_BO_3_, 0.19 g/L of CoCl_2_ ×6H_2_O, 2 mg/L of CuCl_2_ ×2H_2_O, 24 mg/L of NiCl_2_ ×6H_2_O, and 36 mg/L of Na_2_MoO_4_ ×2H_2_O), 5 mL/L of 1M MgCl_2,_ 1 mL/L of 1M CaCl_2_, 100 mL/L of the mixed amino acids and nucleic acids stock (1.6 g/L of alanine, 1 g/L of arginine, 0.4 g/L of asparagine, 2 g/L of aspartic acid, 0.05 g/L of cysteine, 6 g/L of glutamic acid, 0.12 g/L of glutamine, 0.8 g/L of glycine, 1 g/L of histidine monohydrochloride monohydrate, 2 g/L of isoleucine, 2.6 g/L of leucine, 2.4 g/L of lysine monohydrochloride, 0.6 g/L of methionine, 2 g/L of phenylalanine, 2 g/L of proline, 1 g/L of serine, 0.7 g/L of threonine, 0.3 g/L of tryptophan, 0.25 g/L of tyrosine, 2 g/L of valine, 2 g/L of adenine hemisulfate salt, and 2 g/L of uracil), 1.10 mg/L of FeSO_4_ ×7H_2_O, 0.6% final dextrose, and 1% Tween-80.

### Supernatant assay

CFSMs were prepared by inoculating 12 mL of 10%THY+T or BAAD+T medium with cells at an initial OD_600_ of 0.01 (~8×10^6^ CFUs/mL). Cultures were grown for ~16–18 h at 37°C (with shaking), allowing all isolates to reach stationary phase. CFSM was isolated by centrifuging down the cells (3,500 rpm for 10 min) and filtering the supernatant through a 0.22-µm syringe filter. The pH of the CFSM was then re-adjusted (to pH 7.2).

### PRISM assay setup

CFSMs were prepared similar to the supernatant assay described above. The 96 well plate was set up with a different ratio of CFSM to fresh medium (either 10%THY+T or BAAD+T) in a range of 16 µL, 25 µL, 50 µL, 75 µL, 100 µL and 125 µL of CFSM from column B through column G and the opposite direction for fresh media. Additionally, each row from row 1 to row 3 also had a different dilution of the fresh media calculated based on the maximum linear growth of each strain.

The OD_600_ of each well was read every 10 min for 24h using a BioTek Epoch 2 microplate reader. Carrying capacity (using maximum OD_600_ as a proxy) were calculated using code developed in MATLAB R2023. Carrying capacities are estimated based on the maximum OD_600_ values reached for monocultures within 24 h of growth.

### Error estimation and statistics

We modeled the relationship between the growth of a strain to its dependency on resources and mediators using a linear model with a normal (gaussian) distribution. Model parameters were estimated using MATLAB Statistics and Machine Learning Toolbox using the glmfit function. To evaluate the reliability of our estimates, we calculated the Standard Errors for all coefficients and used the model deviance as a measure of goodness of fit.

## Acknowledgements

This work was supported by the National Science Foundation (NSF MCB) under Grant No. 2430384. The authors would like to thank Dr. Katherine P. Lemon for sharing the strains and offering general guidance.

## Code availability

All the relevant codes are available on GitHub at https://github.com/warrierv-cloud/PRISM-codes-2026.git

## Supplemental Material

### S.1 Correcting for the effect of residual resources in the CFSM assay

Cell-free conditioned media (CFCM) are different from cell-free spent media (CFSM) in that the nutrients in the environment are not completely depleted by the effector species. The amount of remaining nutrients in the CFCM is difficult to control, leading to inconsistent estimates of interaction coefficients. Assuming an LV model, the presence of another species modulates the growth rate proportionally to the size of the interacting partner, i.e.

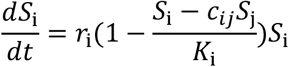

Growth of species *i* in the CFCM of species *j* can be represented as

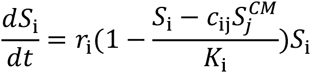

where 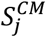 is the density of population *j* at the time of its sampling to obtain the CFCM. This is based on the assumption that in the CFCM of species *j*, the environment will resemble the situation at which the density of *S*_*j*_ has reached 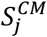.

The carrying capacity of species *i* in CFCM of species *j* is reached when the population *S*_*i*_ reaches a level at which its growth rate becomes zero, thus

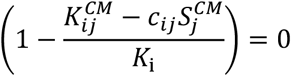

Therefore, the carrying capacity in the CFSM assay is

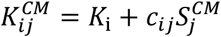

where *K*_i_ is the carrying capacity of species *i* in a monoculture. This allows us to estimate the interaction coefficient *c*_ij_ as

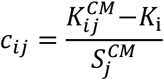

To estimate 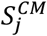, we examine the growth of species *j* in its own CFCM:

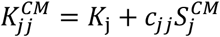

However, we know that in the ideal case, *c*_*jj*_ = −1 (i.e. each species occupies the same niche as itself). Therefore,

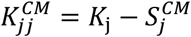

And thus

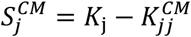

Using this in our estimate of the interaction coefficient, we find

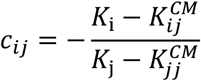

### S.2 Identifying suitable nutrient ranges for linear modeling

To determine the conditions under which our linear regression PRISM model could be applied, we first tested the linear correlation between the bacterial growth rates and carrying capacities over a range of fresh medium dilutions. In highly diluted media (10%THY or more diluted), we see a linear correlation between growth rates and carrying capacities across the three replicates [10]. For four out of the six strains tested (*C. accolens, C. tuberculostearicum, D. pigrum* and *C. pseudodiptheriticum*) we see this linear trend up to 50% THH medium. In more concentrated media, the carrying capacities proportionally increased, but the growth rates deviated from the linear correlation. We posit that in such concentrated media the linear model may no longer be used. The *Staphylococcus* species (*S. aureus* and *S. epidermidis*) showed a linear trend even at higher concentrations of fresh media. For the purpose of partitioning microbial interactions, we limited our investigations to the linear window of medium concentrations to avoid the effects of nutrient saturation.

### S.3 The PRISM assay accurately represents the impact of a known compound on the interaction between microbes

To validate that our linear RM model produces realistic inferences, we tested if the influence of a known compound such as an antibiotic can be inferred from our PRISM assay. For this, we first measured the growth of *S. aureus* and *C. accolens* in response to a known antibiotic compound, ceftazidime. We observed strong inhibition of *S. aureus*, but only a modest inhibitory effect on *C. accolens*. We then added ceftazidime at a concentration of 20 µg/mL to our media, simulating the condition in which another organism has produced this antibiotic compound as an interaction mediator. The media with added antibiotic was treated as the supernatant in this case while setting up the PRISM assay. We observe that this antibiotic as the interaction mediator has an elevated effect on *S. aureus* (final OD_600_ in batch culture was 0.7 lower than the no-ceftazidime control) in comparison to *C. accolens* (final OD_600_ in batch culture was 0.1 lower than the no-ceftazidime control) as represented in Figure S1. We also compared the growth curves of both the strains in the presence and absence of antibiotics. We saw similar results for the effect of ceftazidime on *S. aureus* and *C. accolens* drop in growth as observed from our PRISM assay. This verifies that the PRISM assay correctly represents the effect of a known compound present in CFCMs.

**Figure S1.**
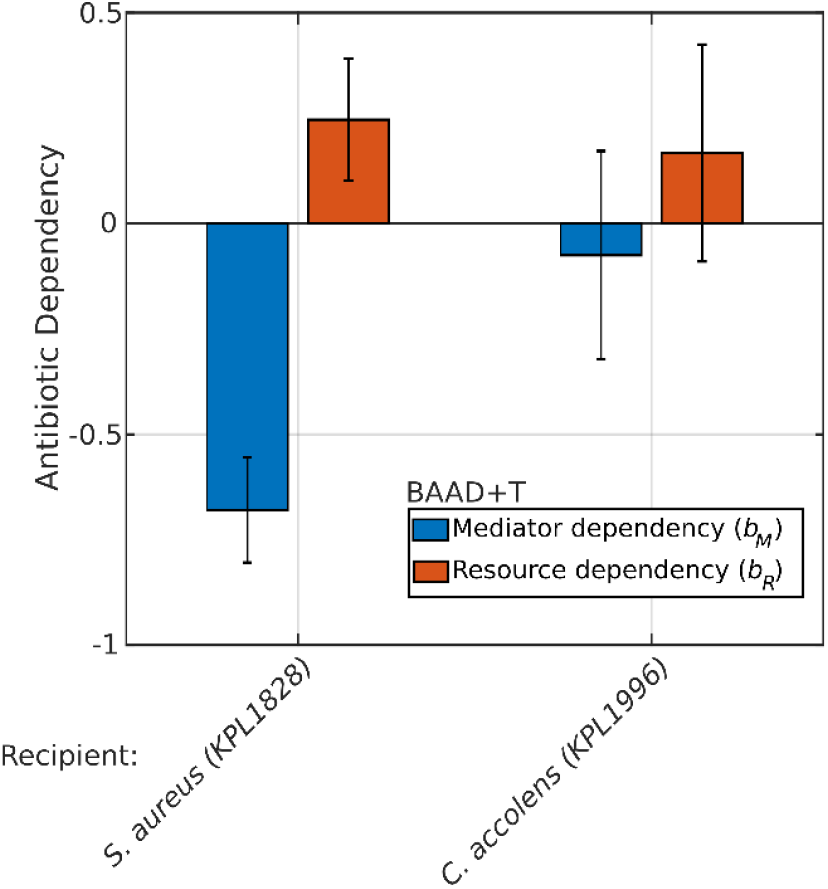
Testing for a known inhibitor such as antibiotic ceftazidime shows that the effect of ceftazidime is more prominent on *S. aureus* than *C. accolens*. The effect of 20 µg/mL of ceftazidime used instead of the supernatant in the PRISM assay is shown. The orange bars represent the dependency of growth on fresh medium, and the blue bars represent the dependency of the growth on the known antibiotic inhibitor ceftazidime. Two strains are tested, *S. aureus* (more sensitive to ceftazidime) and *C. accolens* (less sensitive to ceftazidime). Error bars show the 95% confidence interval obtained from the linear regression of slopes.

